# The SARS-CoV-2 ORF10 is not essential *in vitro* or *in vivo* in humans

**DOI:** 10.1101/2020.08.29.257360

**Authors:** Katarzyna Pancer, Aleksandra Milewska, Katarzyna Owczarek, Agnieszka Dabrowska, Wojciech Branicki, Marek Sanak, Krzysztof Pyrc

**Author notes:** equal contribution (Katarzyna Pancer, Aleksandra Milewska). Correspondence should be addressed to **Krzysztof Pyrc** or **Katarzyna Pancer** or **Marek Sanak** www: http://virogenetics.info/.

## Abstract

SARS-CoV-2 genome annotation revealed the presence of 10 open reading frames (ORFs), of which the last one (ORF10) is positioned downstream the N gene. It is a hypothetical gene, which was speculated to encode a 38 aa protein. This hypothetical protein does not share sequence similarity with any other known protein and cannot be associated with a function. While the role of this ORF10 was proposed, there is a growing evidence showing that the ORF10 is not a coding region.

Here, we identified SARS-CoV-2 variants in which the ORF10 gene was prematurely terminated. The disease was not attenuated, and the transmissibility between humans was not hampered. Also *in vitro*, the strains replicated similarly, as the related viruses with the intact ORF10. Altogether, based on clinical observation and laboratory analyses, it appears that the ORF10 protein is not essential in humans. This observation further proves that the ORF10 should not be treated as the protein-coding gene, and the genome annotations should be amended.

## Introduction

Coronaviruses are mammalian and avian RNA viruses, with large genomes of ~30,000 bases, which encode several proteins required for the virus replication, modulating the immune responses, and forming the scaffold of progeny virions^1^. The spatial distribution of the open reading frames (ORFs) is similar across the taxa. The 1a/1ab ORF starts near the 5’ terminus and is the only ORF that may be translated directly from the genomic RNA, giving rise to the non-structural proteins that re-shape the cellular microenvironment and initiate the replication process. Downstream, a number of ORFs encoding the structural proteins are located (HE, S, M, E, N), interspaced with genes encoding accessory proteins, varying in number and position^1^. SARS-CoV-2 genome annotation revealed the presence of 10 ORFs, of which the last one (ORF10) is positioned downstream the N gene^2^. It is a hypothetical, 117 nt - long ORF, which was speculated to encode a 38 aa protein^2,3^. Bioinformatic analyses revealed that this hypothetical protein does not share the sequence similarity with any other known protein, and the predicted structure cannot be associated with a function. Nonetheless, it was speculated that the ORF 10 protein may play a role in the immunogenicity of the SARS-CoV-2 or may modulate the virulence of the SARS-CoV-2. On the other hand, there is growing evidence showing that the ORF10 is not a coding region. Jungreis *et al*. analyzed the region for different *Sarbecoviruses* and found that only in a minority of cases, for the closest SARS-CoV-2 relatives, the ORF10 is intact. The evidence for the presence of the subgenomic mRNAs corresponding to the ORF10 is limited^4,5^.

Here, we identified two patients infected with the SARS-CoV-2 virus, in which the ORF10 gene was prematurely terminated with a stop codon. The disease was not attenuated, and the transmissibility was not hampered. Isolation of these viruses in cell culture showed that also *in vitro*, these strains replicated similarly, as the related viruses with the intact ORF10.

Altogether, based on clinical observation and laboratory analyses, it appears that the ORF10 protein is not essential in humans.

## Materials and Methods

### Cells and the virus

Vero E6 (*Cercopithecus aethiops*; kidney epithelial; CRL-1586) were cultured in Dulbecco’s MEM (Thermo Fisher Scientific, Poland) supplemented with 3% fetal bovine serum (heat-inactivated; Thermo Fisher Scientific, Poland) and antibiotics: penicillin (100 U/ml), streptomycin (100 μg/ml), and ciprofloxacin (5 μg/ml). Cells were maintained at 37°C under 5% CO_2_.

The strains with the nonsense mutation in the ORF10 gene were designated names PL_P32 and PL_P33 [GISAID Clade G, Pangolin lineage B.1] (accession numbers for the GISAID database: hCoV-19/Poland/PL_P32/2020 and hCoV-19/Poland/PL_P33/2020, respectively) and the reference samples showing high similarity on the nucleotide level, but lacking the point mutation, were designated names PL_P31 [GISAID Clade G, Pangolin lineage B.1] and PL_P38 [GISAID Clade G, Pangolin lineage B.1.5] (accession numbers for the GISAID database: hCoV-19/Poland/PL_P31/2020 and hCoV-19/Poland/PL_P38/2020). Reference SARS-CoV-2 strain 026V-03883 was kindly granted by Christian Drosten, Charité – Universitätsmedizin Berlin, Germany by the European Virus Archive - Global (EVAg); https://www.european-virus-archive.com/).

All SARS-CoV-2 stocks were generated by infecting monolayers of Vero E6 cells. The virus-containing liquid was collected at day 3 post-infection (p.i.), aliquoted and stored at −80°C. Control samples from mock-infected cells was prepared in the same manner. Virus yield was assessed by titration on fully confluent Vero E6 cells in 96-well plates, according to the method of Reed and Muench. Plates were incubated at 37°C for three days, and the cytopathic effect (CPE) was scored by observation under an inverted microscope.

### Sequencing

Total RNA was isolated from the throat swabs collections stored as frozen PBS suspensions at −20°C using a manual TRI Reagent – chloroform extraction and sodium acetate – ethanol precipitation (Sigma-Aldrich, Poznań, Poland). Presence of SARS-CoV-2 material in the collected sample was tested using GeneFinder real-time COVID-19 plus kit (OSANG Healthcare, Korea). Isolated total RNA was treated with DNAse I to remove DNA contamination, reverse transcribed with SuperScript IV and random oligohehamer primers, next second strand synthesis was compleded using DNA polymerase I (all reagents from Thermo Fisher, Warszawa, Poland). Illumina platform sequencing libraries were prepared using Nextera Flex Enrichment Library with Respiratory Virus Oligo Panel capture worflow according to the manufacturer instruction Illumina – Analitik, Warszawa, Poland). Two libraries of 12 samples barcoded with individual i7 and i5 adapters were sequenced in each run. NGS sequencing was accomplished using MiSeq v.3 2×75 chemistry (Illumina). Raw sequencing files were demultiplexed using IlluminaBasecallsToFasq procedure from PICARD package and mapped to NC_055512.2 SARS-CoV-2 reference sequence with BwaAndMarkDuplicatesPipelineSpark procedure from GATK v.4.1.5.0 package (Broad Institute, Boston, MA). Individual samples files were manually inspected using Integrated Genomics Viewer (Broad Institute). Only 2 samples out of 72 sequenced had identical C>T transition at NC_0055512:29642 position within the putative orf10 at 3’ of the virus genome. Base T read quality value was QV=38 and numbers of reads were 265 and 340 for samples PL_P32 and PL_P33. This transition could change putatione codon 29 from glutanine (CAA, id-gu280_gp11.2) to the stop (TAA). No other sequence variants were detected in the orf10 region. Sequence alignments of samples PL_P32 and PL_P33 are in the **Supplementary File 1**.

### Isolation of nucleic acids and reverse transcription

A viral DNA/RNA kit (A&A Biotechnology, Poland) was used for nucleic acid isolation from cell culture supernatants. RNA was isolated according to the manufacturer’s instructions. cDNA samples were prepared with a high-capacity cDNA reverse transcription kit (Thermo Fisher Scientific, Poland), according to the manufacturer’s instructions.

### Quantitative PCR

Viral RNA was quantified using quantitative PCR (qPCR; CFX96 Touch real-time PCR detection system; Bio-Rad, Poland). cDNA was amplified using 1× qPCR master mix (A&A Biotechnology, Poland), in the presence of the probe (100 nM; FAM/BHQ1, ACT TCC TCA AGG AAC AAC ATT GCC A) and primers (450 nM each; CAC ATT GGC ACC CGC AAT C and GAG GAA CGA GAA GAG GCT TG). The heating scheme was as follows: 2 min at 50°C and 10 min at 92°C, followed by 30 cycles of 15 s at 92°C and 1 min at 60°C. In order to assess the copy number for the N gene, standards were prepared and serially diluted.

## Results and Discussion

The first SARS-CoV-2 infected patient was identified in Poland on the 4^th^ of March 2020, and the subsequent monitoring of the genetic drift of the virus was initiated and allowed for the characterization of circulating viruses. The phylogenetic analysis led to the conclusion that the diversity of the virus is similar to the one observed worldwide^6^. The virus was introduced to the population of Poland from different sources, as hallmarks of different clades are present; virtually all genetic clades identified thus far were present^6^. In the course of analysis, some isolates showed some peculiarities. In two samples, sequencing revealed the disruption of the ORF10, as a stop codon was present at position aa 29. This premature termination results from the C-T mutation, amending the CAA to TAA codon, what is characteristic for coronaviral genomes^7,8^. The sequence of this particular region was covered >260 times, and no minority variants were detected. Interestingly, our submission of the sequence to the GenBank database was rejected, due to the stop codon in the ORF10 gene.

As we already knew that both original samples carried this mutation, we analyzed the accessible clinical data. A 58 year-old Polish man living in Warsaw, Poland, spent a few days in the Germany at the end of the February, 2020. After returning to Poland, he was informed that he was in contact with the person infected with the SARS-CoV-2 virus. Despite the lack of obvious symptoms, he contacted a public health center. On the 4^th^ day from the exposure (the 5 ^th^ March, 2020) throat swab was collected and transported in saline medium. The same day, real-time RT-PCR analysis was carried out in the National Institute of Public Health – NIPH, in Warsaw using the Primerdesign, genesig Real-Time PCR CoVID-19 kit. RT-qPCR according to Charite protocol was used for verification of the result^9^. The sample was stored and sequenced (hCoV-19/Poland/PL_P32/2020). In due course, a symptomatic infection developed and the patient experienced fatigue and loss of smell and taste. No symptoms of the respiratory tract infection were reported. The infection lasted for 26 days, and the person recovered with no sequelae. On the 6^th^ of March, 2020, the patient’s wife (female, 62 years) was examined and the result was inconclusive. However, the second sample collected on 13^th^ of March, 2020 was positive for the SARS-CoV-2 RNA. The sample was stored and sequenced (hCoV-19/Poland/PL_P33/2020). The patient experienced fever (38°C) for 2 days and recovered without sequelae. Further, the sample was also collected from another person that was in contact with the first case (male, 41 years). Sample was tested positive for the SARS-CoV-2 RNA. No sample was collected for sequencing. The patient experienced a dry cough and loss of smell and taste. The infection lasted for 21 days, and the person recovered with no sequelae.

Based on the collected data, one may safely assume that the virus with the disrupted ORF10 was infectious and pathogenic in humans. The identical change in two patients proves that it was not resulting from intra-patient genetic drift and that the virus transmissibility was not affected.

To further characterize the phenotype of the virus, available clinical samples were overlaid on the fully confluent Vero E6 cells. At the same time, parallel cultures were inoculated with closely related PL_P31 and PL_P38 isolates (see **Figure 1**).

**Figure 1.**
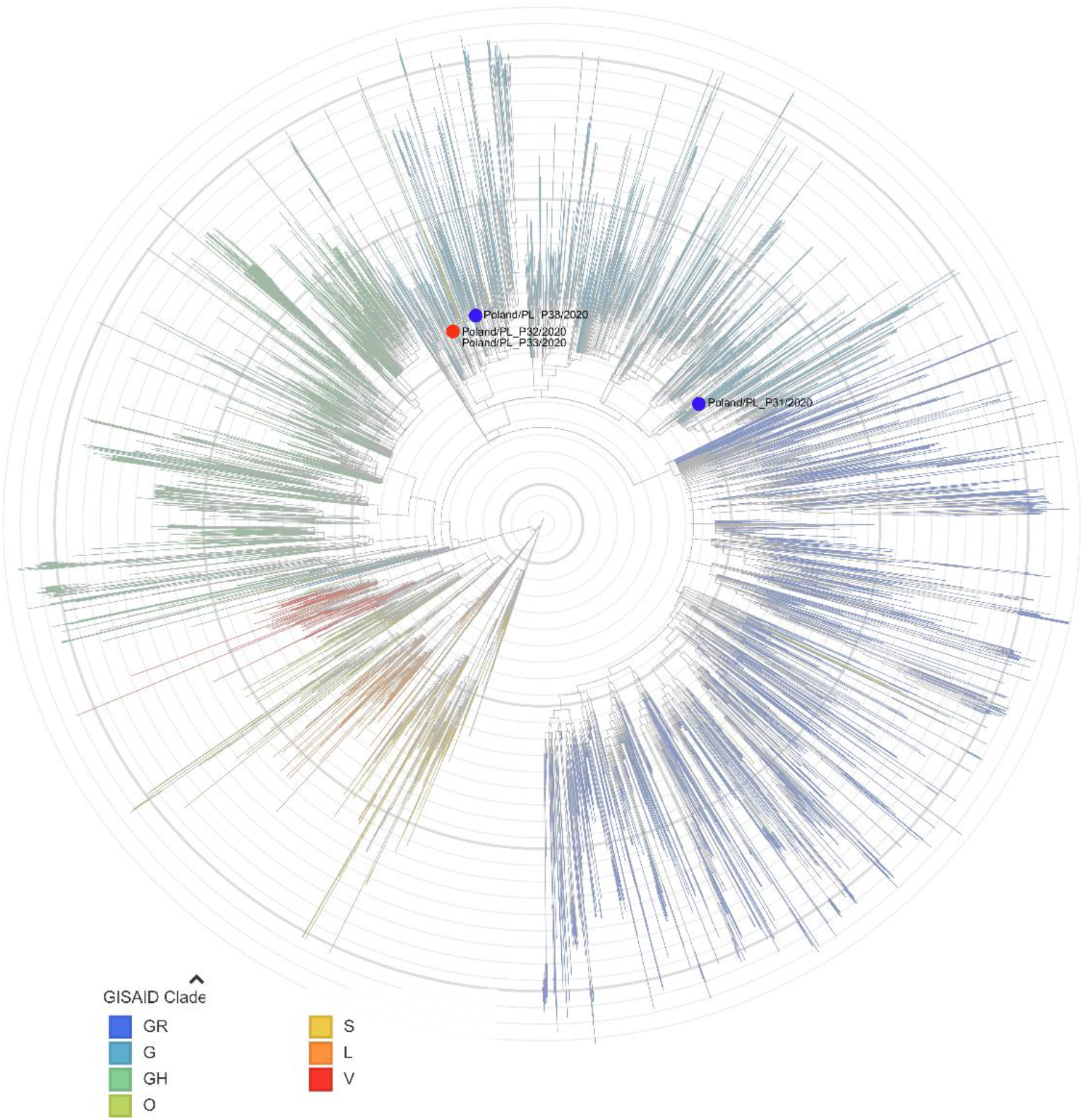
Phylogenetic analysis of the isolates included in the study. The analysis was carried out using the nexstrain server, with the dataset dated on 4^th^ of August 2020^10,11^. The strains with the point mutation in the ORF10 are labeled in red, while the reference strains are labeled in blue.

In all four cases, 72 h post-inoculation we observed the appearance of characteristic CPE. The media samples were collected daily, and total RNA was isolated. The RT-qPCR reaction was carried out, and the virus yields are presented in **Figure 2**. No difference between the replication dynamics between strains carrying the nonsense mutation in the ORF10 and the strains with intact ORF10 was observed.

**Figure 2.**
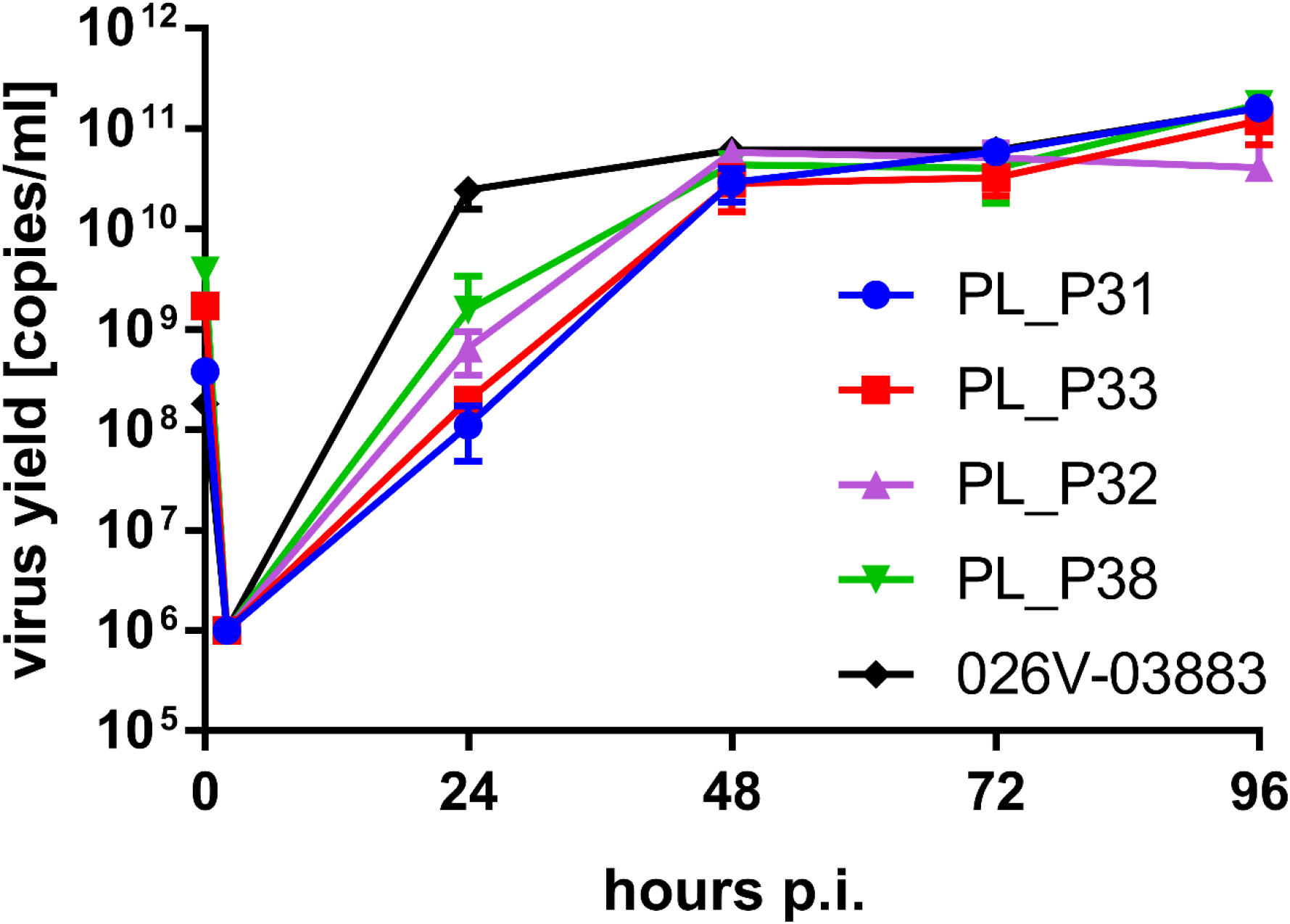
Replication kinetics of the SARS-CoV-2 strains. Virus yield wass determined with RT-qPCR and the data is presented as a mean ±SD. The EVAg strain was used as a reference.

Concluding, results obtained from the cell culture, sequencing, and clinical data show that the stop codon in the two-thirds of the protein did not affect the virus fitness. This observation further supports the thesis that the ORF10 should not be treated as the protein-coding gene, and the genome annotations should be altered^4^. On the other hand, ORF10 is relatively conserved, suggesting the importance of this region, e.g., due to the secondary RNA structures.

## ACKNOWLEDGEMENTS

This work was supported by the subsidy from the Polish Ministry of Science and Higher Education for the research on the SARS-CoV-2 and a grant from the National Science Center UMO-2017/27/B/NZ6/02488 to KP.

Authors thank Illumina Netherlands BV for the consumables including Respiratory Virus Oligo Panel, provided free of charge in connection with exploring research and surveillance in response to the SARS CoV-2 pandemic.

The funders had no role in study design, data collection, and analysis, decision to publish, or preparation of the manuscript.

The authors declare no competing financial interests.

